# Allelic variation in MAL33 drives ecological adaptation of maltose metabolism in Saccharomyces eubayanus

**DOI:** 10.1101/2025.09.15.676268

**Authors:** Pablo Quintrel, Felipe Muñoz-Guzmán, Pablo Villarreal, Tomas A. Peña, Natalia I. Garate, Catalina Muñoz-Tapia, Christian I. Oporto, Johnathan G. Crandall, Luis F. Larrondo, Chris Todd Hittinger, Gilles Fischer, Francisco A. Cubillos

## Abstract

Maltose is one of the most abundant sugars in brewer’s wort, and its efficient utilization is critical for successful fermentation. However, maltose consumption varies naturally among *Saccharomyces eubayanus* strains isolated from different host trees, such as *Quercus* and *Nothofagus*. To identify the genetic determinants underlying these phenotypic differences, we performed bulk segregant analysis (BSA) and quantitative trait loci (QTL) mapping using an F2 offspring derived from QC18 (Quercus-associated) and CL467.1 (Nothofagus-associated) strains. QTL mapping identified two significant genomic regions on subtelomeric loci of chromosomes V-R and XVI-L, each containing complete *MAL* loci composed of *MAL32* (encoding maltase), *MAL31* (transporter), and *MAL33* (transcriptional activator) genes. Comparative polymorphism analyses identified mutations in *MAL32* and *MAL33* of QC18, including frameshift mutations resulting in premature stop codons. Functional validation demonstrated that the heterologous expression of *MAL33*_ChrV_ from CL467.1 fully restored maltose utilization in QC18, indicating the functional presence of *MAL33* cis-regulatory sequences and *MAL32* and *MAL31* genes in QC18. While structural protein predictions identified truncation and impaired functionality in the maltose-responsive activation domain of Mal33p from QC18, overexpression of QC18’s own *MAL33*_ChrV_ allele also improved maltose metabolism, suggesting dosage-dependent transcriptional limitations rather than complete functional loss. These results indicate that allelic variations in the maltose-responsive activation domain of Mal33p lead to differences in maltose consumption between strains. We hypothesized that reduced maltose metabolism in QC18 is an adaptive response to the distinct sugar composition in *Quercus robur* bark, contrasting with the starch-rich environment of *Nothofagus pumilio*. These findings highlight subtelomeric *MAL* gene diversity as a reservoir of evolutionary plasticity, representing a key evolutionary mechanism that influences maltose adaptation among natural *Saccharomyces* isolates.

## Introduction

Microorganisms frequently encounter environmental fluctuations that challenge their fitness (Jacob & Monod, 1961; Perez-Samper et al., 2018). To adapt, they rely on gene regulatory circuits that integrate extracellular signals and trigger rapid responses (New et al., 2014; Rashida et al., 2021; Ricci-Tam et al., 2021). A classic example is diauxic growth, where cells sequentially utilize carbon sources (Bheda, 2020). In the yeast *Saccharomyces cerevisiae*, glucose is the preferred carbon source and is transported into cells via diffusion carriers encoded by hexose transporter genes (*HXT*) (Hatanaka et al., 2018, 2022). Glucose represses the expression of gene families associated with other non-preferred sugars such as *MAL*, *GAL*, or *SUC* (for maltose, galactose, and sucrose consumption, respectively) (Fita-Torró et al., 2023; Harrison et al., 2022, 2022; Kayikci & Nielsen, 2015; Mao & Chen, 2019; Neigeborn & Carlson, 1984; New et al., 2014; Perez-Samper et al., 2018; Stewart et al., 2013; Stockwell et al., 2015). The Snf1p/Mig1p proteins mediate this repression (Fita-Torró et al., 2023), ensuring that metabolic resources are exclusively dedicated to optimal glucose consumption (Perez-Samper et al., 2018). During the diauxic lag phase, several genes undergo derepression and subsequently activation to synthesize the necessary enzymes for utilizing non-preferred sugars, thereby resuming exponential growth (New et al., 2014; Venturelli et al., 2015). This metabolic reprogramming can persist for several hours, depending on the yeast strain (J. Wang et al., 2015), creating a hiatus before exponential growth resumes, a delay that can be detrimental to industrial processes if extended. For example, maltose is the primary carbon source in beer fermentation (He et al., 2014), and yeast cells must rapidly switch from glucose to maltose metabolism. Failure to do so during beer production (traditionally referred to as stuck or sluggish fermentations) can result in a sweet, spoilage-prone final product, leading to significant economic losses (Perez-Samper et al., 2018; Verstrepen et al., 2004).

The *MAL* locus bears the essential genetic information for maltose utilization (Brickwedde et al., 2018; Needleman & Michels, 1983). It consists of a transcriptional activator that controls the expression of a maltose permease and alpha-glucosidase (maltase) encoded by *MALx3*, *MALx1*, *and MALx2,* respectively (Chow et al., 1989; Naumov et al., 1994). In *S. cerevisiae*, the *MAL* locus is commonly found in subtelomeric regions, and exhibits copy number variation among strains, which impacts their ability to use maltose (Brown et al., 2010; Duval et al., 2010; Peter et al., 2018). Effectively consuming and utilizing maltose is considered a hallmark of domestication in *S. cerevisiae* and a desirable trait in brewing (Duval et al., 2010; Gallone et al., 2016, 2018; Molinet et al., 2022). *S. cerevisiae* strains from different populations exhibit a broad spectrum of diauxic lag phenotypes and gene expression changes related to maltose consumption (Caudal et al., 2024; Gonçalves et al., 2016; Hernández-Vásquez et al., 2024; Warringer et al., 2011). Copy number variation in the *MAL* locus, particularly in the maltose transporter genes, also contributes to this enhanced performance (Gallone et al., 2016; Gonçalves et al., 2016). Although the *MAL* locus is typically organized in a functionally conserved locus across the *Saccharomyces* genus (Brouwers et al., 2019), *Saccharomyces* yeast strains isolated from wild environments lack this domestication signature and typically exhibit slower maltose consumption rates and longer diauxic lag phases than commercial beer strains (Gonçalves et al., 2016; Hutzler et al., 2021; Molinet et al., 2022; Nikulin et al., 2020). For example, natural isolates of *Saccharomyces eubayanus*, the cold-tolerant parental species of the industrial hybrid *Saccharomyces pastorianus*, tend to exhibit a slow diauxic lag phase under wort conditions, likely due to strong glucose repression (Mardones et al., 2022).

The ability to consume maltose is present across most sub-populations of *S. eubayanus*. Wild *S. eubayanus* isolates from Patagonia have two complete copies of the *MAL* locus (Brickwedde et al., 2018; Brouwers et al., 2019) and exhibit wide phenotypic variation in maltose growth and consumption leading to differences in their fermentative capacity (Molinet et al., 2022; Nespolo et al., 2020). However, maltose consumption is rather absent in the Holarctic strains, which are hypothesized to be lager yeast’s direct parental lineage (Bergin et al., 2022). This may be attributed to an altered maltose expression profile in the *MAL* locus (Baker et al., 2019; Crandall et al., 2023; Eizaguirre et al., 2018; Langdon et al., 2020). Most notably, Holarctic yeasts in the Northern Hemisphere have only been isolated from *Quercus robur*, including *S. eubayanus* isolates from Ireland (Bergin et al., 2022). Therefore, it is relevant to highlight that the QC18 strain isolated from an exotic *Q. robur* tree in Argentina and belonging to the Patagonian A lineage also displayed a low fermentative profile, poor maltose consumption, and a slow adaptation to maltose in mixed media containing glucose, which can last several days (Molinet et al., 2022). Gene expression profiling revealed *CIN5* allelic variants impacting diauxic shift between *S. eubayanus* strains. However, these variants did not recapitulate their phenotypic differences, suggesting additional genetic determinants underlying the phenotypic differences between strains. In this sense, the genetic determinants underlying maltose consumption and gene expression differences across *S. eubayanus* strains remain unknown.

To elucidate the genetic basis of maltose consumption variation in *S. eubayanus*, we investigated two strains exhibiting contrasting phenotypes in beer wort fermentation and maltose consumption: CL4671.1 and QC18, representing a short and long lag phase during diauxic shift experiments, respectively. We characterized their phenotypic response under various conditions using quantitative trait loci analysis to uncover the genetic factors responsible for maltose consumption during fermentation. We implemented a complementation assay, which restored the QC18 maltose consumption trait, providing insight into the regulation of maltose consumption. Additionally, we characterized the phenotypic contribution of the two *MAL* locus copies in QC18 and CL467.1 strains, demonstrating a differential contribution to the maltose consumption phenotype. These results highlight the evolutionary role of gene families in subtelomeric regions and provide a plausible ecological explanation for the natural variation in maltose consumption observed among natural *S. eubayanus* strains.

## METHODS

### Strains and storage media

The strains used in this work are listed in **Table S1** (Molinet et al., 2022). All the strains were maintained on YPD solid media (1% yeast extract, 2% peptone, 2% glucose, 2 % agar). For long-term storage, all strains were frozen in storage media (YPD, glycerol 20%) at −80°C.

### Diauxic growth phenotypic characterization

Strains were phenotypically characterized in microculture conditions in 96-well plates under different diauxic media conditions. Carbon sources and conditions are listed in **Table S2**. Pre-cultures were grown 24 hours on YP (1% yeast extract, 2% peptone) supplemented with 5% glucose as the only carbon source for 24 hours at 25°C. Then, the pre-culture was refreshed by diluting 20 μL of the initial culture with 180 μL of YP 5% glucose for 24 hours at 25°C. On the third day, 20 μL of the pre-culture were washed in 180 μL of YP media, and then 20 μL were inoculated to the media array of different carbon sources to a final volume of 200 µL. For the main experiment, we denominated ‘sudden’ and ‘gradual’ diauxic shift experiments if cells were incubated in a single carbon source or a mixture of carbon sources, respectively. Growth was constantly monitored every 60 minutes at OD_620_ in a TECAN sunrise instrument at 20°C. For the large-scale segregant pool phenotyping, plates were incubated on a SPECTROstar Omega plate reader (BMG Labtech) equipped with a microplate stacker, and OD_620_ was measured every hour. Biomass generation was quantified using the Area Under Curve (AUC), using the GrowthCurver package built under R software version 4.1.2. All growth measurements represent the averages of at least three biological replicates.

### Generation of F1 Hybrids and F_2_ segregants between CL467.1 and QC18

The CL467.1 and QC18 hybrid was constructed using a wild-type version of QC18 and a stable haploid version of CL467.1 (Molinet et al., 2022). For this, the QC18 strain was sporulated on 2% potassium acetate agar plates (2% agar) for at least seven days at 12 °C. Crosses were performed between a QC18 spore and an isolated CL4671.1 cell using a SporePlay micromanipulator (Singer, UK). Crosses were checked by colony PCR using the *HO* gene as a marker as previously described (Molinet et al., 2022). All the primers are listed in **Table S3**.

F_2_ segregants were obtained by sporulating the F_1_ CL4671.1 x QC18 Hybrid, as previously mentioned. Haploid segregants were selected by colony PCR of the *MAT* and *HO* loci (**Table S3**). A total of 273 haploids segregants were obtained and stored for future experiments. An F_2_ segregant is considered transgressive if its total growth value (Area Under the Curve, AUC) is either lower or higher than the phenotypic value of the parental strain QC18 or CL467.1, respectively

### DNA extraction

DNA was obtained from an overnight 100 μL saturated culture for each segregant. Cells were treated with 10uL Zymolyase 100T (50mg/mL) at 37°C for 1 hour, and DNA extraction was performed using the Quick-DNA, Miniprep Kit (ZymoResearch). DNA concentrations were measured in Qubit (Invitrogen). 100 ng of DNA from each segregant were mixed, and two top and bottom DNA pools were generated. Whole genome sequencing of parental strains and segregant pools was performed using Illumina pair-end sequencing (Illumina NextSeq500, Universidad de Santiago de Chile).

### Bioinformatic and QTL mapping analyses

Initially, reads were processed using fastp 0.23.2 (-q 20 -F 15 -f 15) {S. Chen, Zhou, Chen, & Gu, 2018}. Trimmed reads were aligned against the *S. eubayanus* CBS12357^T^ reference genome {Libkind et al, 2011} using BWA-mem (option: -M -R) {Li, 2013}. Mapping quality, summary statistics and variant calling were performed as previously described {Peña et al, 2024}. Specifically, variants were called per chromosome per sample using HaplotypeCaller (default settings). We only considered SNPs without missing data using –max-missing 1 {https://github.com/broadinstitute/gatk}. Allele counts from CL4671.1 and QC18 at each polymorphic site were obtained using bcftools stats command (Danecek et al., 2021). Allele count was carried out using Python 3 language; the Allele frequency graph was plotted using ggplot2 from R software.

LOD scores for allele frequency differences between the pools were calculated using MULTIPOOL in Python 2 Version 0.10.2 (https://github.com/matted-zz/multipool). Allele counts were the input to the mp_inference.pt script in python 2; with the options -m contrast -r 100 -c 2200 -n 30. Regions with LOD score greater than 10 were deemed as QTLs.

### Reciprocal hemizygosity analysis

Null mutants were constructed using CRISPR-Cas9 (Dicarlo et al., 2013) as previously described in *S. eubayanus* {molinet et al, 2024}. Briefly, each gRNA was cloned in the plasmid pAEF5 (Fleiss et al., 2019) using a standard “Golden Gate Assembly” (Horwitz et al., 2015). A KanMX cassette was inserted within the two selected QTL blocks linked to maltose utilization for the Reciprocal hemizygosity analysis. For this, we used, in each case, two gRNAs targeting the block’s extremes and a donor DNA containing a KanMX cassette flanked by 30bp sequences of the target regions. Plasmids and donor DNA were co-transformed using the Lithium acetate protocol (Baker et al., 2019). Correct gene deletion was confirmed by standard colony PCR. All the primers, gRNAs and donor DNA are listed in **Table S3.** Reciprocal hemizygotes were constructed by crossbreeding one wild type strain against another containing the mutated target gene. All the strains generated are listed in **Table S1**.

### Episomal heterologous complementation assay

The *S. cerevisiae* laboratory strain BY4741 was used to assemble the expression plasmid pRS426. A DNA fragment containing overlapping regions with the vector, the *MAL33* ORF, and the immediate 1 kb upstream region of the *MAL33* transcriptional starting site from chromosomes V and XVI from QC18 and CL467.1 strain was amplified by PCR (**Table S3**). The pRS426 plasmid was linearized using the Kpn*I* and EcoR*I* enzymes (NEB, UK) at 37°C for 3 hours, followed by inactivation at 80°C for 20 minutes. The *MAL33* gene constructs and the linearized plasmid were co-transformed into the BY4741 strain using the standard lithium acetate transformation protocol, with uracil as the transformant selection agent. Plasmids were extracted using the Zymoprep yeast miniprep kit II (Zymo Research) and transformed into *DH5-α* competent *Escherichia coli* strains using a standard transformation protocol {Green et al, 2013}. Purified plasmids were transformed into the corresponding *S. eubayanus* strain, using Geneticin (G418) as a selection marker.

### Protein structure prediction

To predict the tertiary structure of the Mal33 regulatory protein from both QC18 and CL467.1 genomes, we utilize the ColabFold notebook, a Google Colab-based AlphaFold2 (Mirdita et al., 2022). Protein sequences translated from FASTA files of *MAL33_ChrV_* and *MAL33_ChrXVI_* from QC18 and CL467.1 strains were used as input. For each protein prediction we use the developer’s best practices recommendation (Kim et al., 2024). Multiple sequence alignments (MSA) were generated using MMseqs2 against UniRef and environmental databases, and template search was enabled to include PDB70-derived templates. The AlphaFold2 pipeline was executed using the five standard pre-trained models, with default settings including three recycling cycles and Amber-based structural relaxation. Each protein generated five models ranked by confidence score pLDDT, and the top-ranked model (rank 1) was selected and exported in PDB format. Finally, the selected rank 1 structure was visualized and rendered using PyMOL.

### Data Accessibility

All sequences have been deposited in the National Center for Biotechnology Information (NCBI) as a Sequence Read Archive under the BioProject accession number PRJNA1314239.

## RESULTS

### Maltose growth differences between S. eubayanus strains

We systematically compared the growth of strains QC18 and CL467.1 at different maltose concentrations. Cells were first grown for 48 hours in YP medium with 5% glucose, then shifted to various maltose levels (ranging from 5% to 30%), and the progression of biomass was measured (Figure 1). Growth was evaluated by calculating the Area Under the Curve (AUC), ODmax, and lag-phase duration after 96 hours (**Figure 1A**). Neither strain showed significant differences in AUC and ODmax at low maltose concentrations (YP-5% maltose, *p*-value > 0.05). However, significant differences in the three analyzed parameters were evident between strains at concentrations above 5% (**Figure 1B**, *p*-value < 0.05). While 10% maltose also revealed distinct behaviors, the most pronounced differences in the three parameters measured occurred in YP-20% maltose. Here, the QC18 strain exhibited AUC and ODmax values that were 5.2x and 2.5x lower than CL467.1 (*p*-value < 0.05, ANOVA), respectively. Additionally, QC18 exhibited a lag phase that was 1.7x more pronounced than CL467.1 (p-value < 0.05, Table S4A), with a lag phase partial exit occurring only after 72 hours (Figure 1C). Moreover, when maltose concentration was increased to 30%, QC18 failed to grow, whereas CL467.1 exhibited only a slight reduction compared to its growth in 20% maltose, albeit a longer lag (Figure 1C).

**Figure 1.**
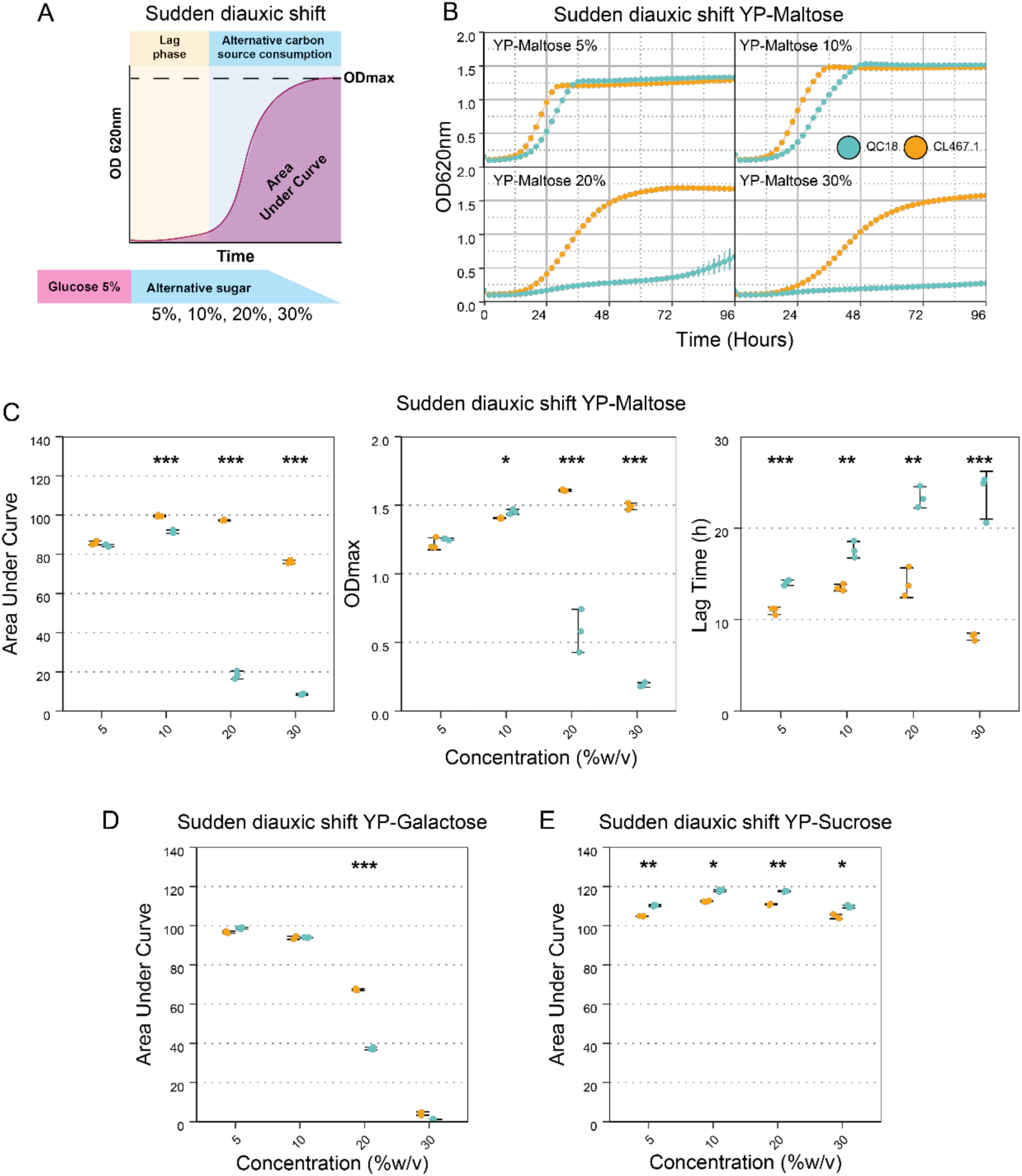
Effect of Alternative Carbon Sources on Sudden Diauxic Growth. **A)** Experimental set-up for sudden diauxic shift experiments screening measuring total growth of native strains of *Saccharomyces eubayanus*. Cultures grown in YPD 5% glucose are transferred to different alternative carbon sources (maltose, galactose or sucrose) at increasing concentrations (5, 10, 20, and 30%; %w/v). **B)** Growth kinetics of QC18 (Turquoise) and CL467.1 (Orange) on YP supplemented with four different maltose concentrations. Lines depict the median OD_620nm_ of three biological replicates. **C**) Three parameters were calculated for maltose growth: total growth was determined by calculating the Area Under Curve (AUC), Lag time (h), and ODmax. QC18 and CL467.1 are shown in Turquoise and Orange, respectively. AUC values were estimated for (**D**) galactose and (**E**) sucrose concentrations. Plotted values correspond to the mean value of three independent replicates for each strain. (*) represents different levels of significance between QC18 and CL467.1 strains in one condition (* *p*-value < 0.05; ANOVA, (**) *p*-value < 0.01 and (***) *p*-value < 0.001.

We also assessed yeast growth under galactose and sucrose (**Figures 1D and 1E**), confirming that the metabolic divergence between QC18 and CL467.1 is exclusive for maltose, and not just sudden shifts to other non-preferred sugars **(Figure S1)**. Across the different sucrose concentrations, we find significant differences in total growth between QC18 and CL467.1 (**Figure 1E**). However, the differences in the three analyzed parameters did not exceed 5%. On the other hand, we found significant differences for 20% galactose (**Figure 1D**), where CL467.1 exhibiting lower AUC values than QC18 (**Table S4**). Despite the significant differences in ODmax (**Figure S1C-D**), both strains reached values above 1.5, suggesting that QC18 can grow on galactose after a sudden shift (**Figure S1D**). These results highlight that the metabolic differences, including an extended diauxic lag phase in QC18, are associated with high levels of maltose, and not just any sugar.

Next, we evaluated the strain’s growth in gradual diauxic shift conditions, using a mixture of YP-1% glucose and different galactose, sucrose, or maltose concentrations. Significant differences were observed when 20% maltose was tested, with the QC18 strain exhibiting a 2-fold lower AUC than CL467.1 (p-value < 0.05). Importantly, these differences stem from QC18 displaying a significantly lower ODmax compared to CL467.1 (**Figure S2A**, p-value < 0.05), rather than from lag phase differences, as seen in the sudden shift assays (**Figure S2A**, p-value > 0.05). showing that the growth of strain CL467.1 is not affected under this condition. Slight differences in AUC were observed between QC18 and CL467.1 in glucose:sucrose and glucose:galactose (**Figure S2B and C**), at concentrations under 30%. At 30%, QC18 exhibited 1.1 times higher growth in sucrose and 1.3 times higher growth in galactose compared to CL467.1. (**Fig S2B and C**, *p*-value > 0.05, ANOVA). These results highlight strain-specific differences, primarily for maltose, in both sudden and gradual diauxic shift conditions.

### QTL mapping reveals two loci associated with maltose utilization differences

To determine the genetic origin of the phenotypic diversity between CL467.1 and QC18 strains, we initially characterized the phenotypic segregation across an F_2_ population (**Table S5**). We generated an F_1_ CL467.1 x QC18 hybrid, and from this, 273 F_2_ haploid segregants were obtained in a single sporulation round (**Figure 2A**). We challenged this F_2_ population to two diauxic shift conditions, sudden in 20% maltose and gradual in 1% glucose with 20% maltose, respectively (**Figure 2B and C**). As a control condition, we evaluated the growth of the segregants in 1% glucose, which showed a narrow distribution of the F_2_ population (**Figure S3**; Coefficient of Variation, CV = 4%), confirming the absence of differential growth under this condition. The sudden condition revealed a broader phenotypic distribution within the segregating population compared to the gradual condition (**Figure 2B**), with a coefficient of variation equal to 41% (**Figure 2B**) and 18% (**Figure 2C**), respectively. In addition, the sudden condition evidenced a significant multimodal distribution with two peaks (**Figure 2D**; D = 0.053, *p*-value< 0.001, Hartigan’s dip test), while the gradual condition exhibited a normal distribution (**Figure 2D**; D = 0.021, *p*-value > 0.05, Hartigan’s dip test). These results suggest that sudden and gradual maltose utilization traits may have distinct genetic determinants, representing quantitative traits with phenotypic segregation within an F_2_ population.

**Figure 2.**
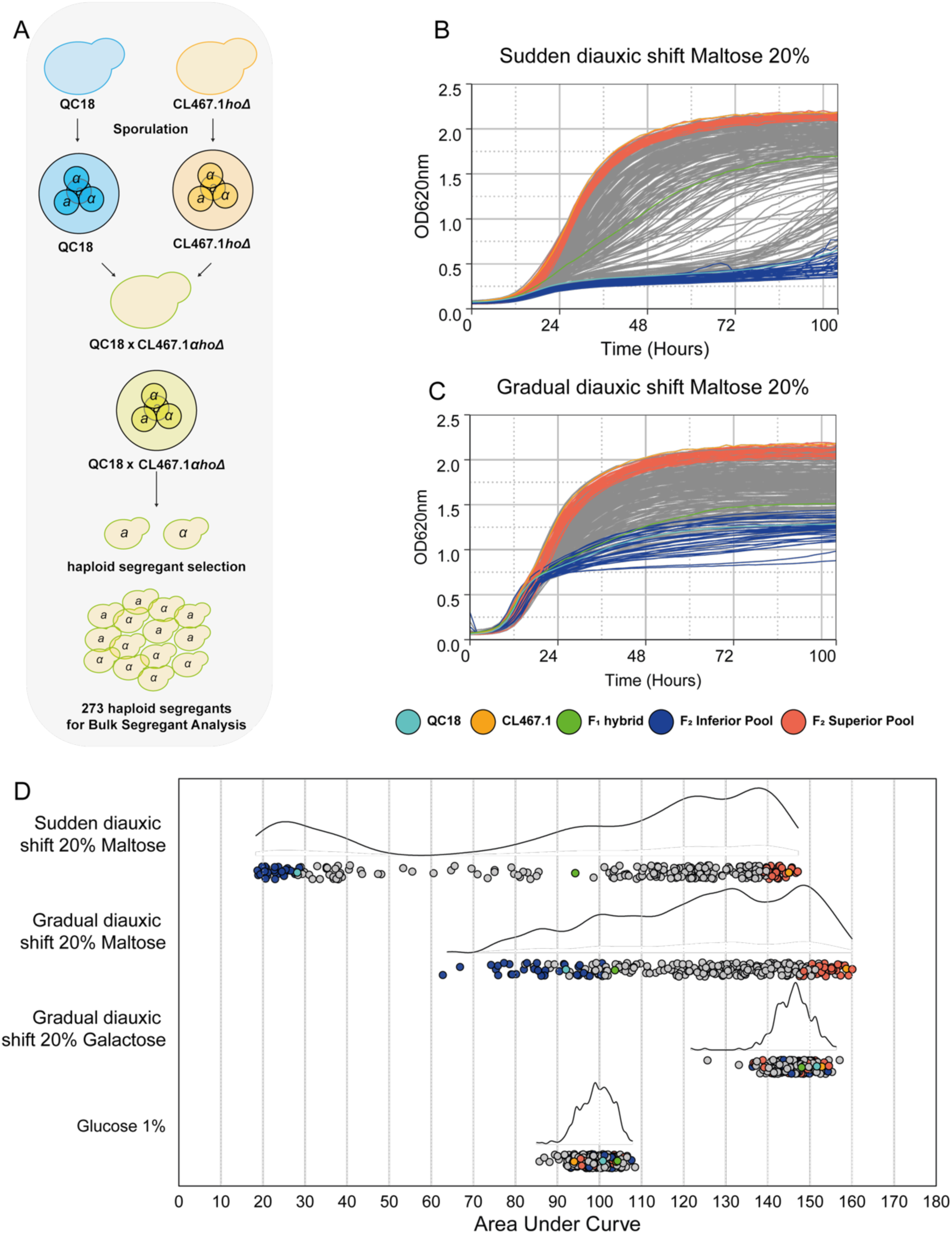
Bulk Segregant Analysis between QC18 and CL467.1 strains under sudden and gradual diauxic shift. **A)** QTL mapping strategy between QC18 and CL467.1. Growth kinetics of QC18, CL467.1, F_1_ hybrid, and F_2_ segregants in (**B**) sudden and (**C**) gradual diauxic shift experiments, using 20% maltose as a secondary carbon source. Parentals strains QC18 and CL467.1 are represented by turquoise and orange lines. Top and bottom 30 F_2_ segregants were selected from each extreme pool, F_2_ Inferior and superior segregants are represented by blue and dark orange lines, respectively. Lines show the OD_620nm_ median for three biological replicates. **D)** F_2_ segregants Area Under Curve (AUC) distribution in diauxic shift experiments, using maltose, galactose or glucose as the secondary carbon source. Each dot represents an F_2_ segregant, turquoise and orange dots represent QC18 and CL467.1 strains; Green represents the F_1_ hybrid; blue and dark orange dots represent the F_2_ Inferior and Superior pools. Dots show the median AUC of three biological replicates.

We chose the sudden diauxic shift strategy for further analysis because of the greater phenotypic variability observed in the F_2_ population. In this way, we identified 3 positive and 31 negative transgressive segregants, respectively (**Figure 2D**). Interestingly, the negative transgressive segregants exhibited a 1% and 53% greater and lower AUC growth relative to the CL467.1 and QC18 strains, respectively (**Figure 2D**).

We employed a QTL mapping approach to identify the genetic variants underlying the differential maltose utilization between the parental strains (**Figure 2D**, Superior and Inferior Pools). For this, we selected 30 Superior and Inferior segregants, subjected them to whole-genome Illumina sequencing, and estimated the allele frequencies for each pool using the CL467.1 genome as a reference (**Figure 3**). We mapped allelic frequency against the genomic position to identify regions with allele frequency differences between both pools and selected regions with LOD scores> 10 (**Table S6**). This analysis led to the detection of two significant QTLs, located in the subtelomeric regions of chromosomes V (QTL_ChrV_) and XVI (QTL_ChrXVI_) **(Figure 3A).** QTL_ChrV_ is in the left arm of chromosome V between 500 kb and 580 kb, (LOD score peak =12.9, Chr5 520kbp, **Figure 3B**). QTL_ChrXVI_ is found in the right arm of chromosome XVI between 10 and 65 kb (LOD score peak = 12.34, ChrXVI 10kbp, **Figure 3C**). In both cases, the top segregants were enriched in the CL467.1 allele, while the bottom segregants were in the QC18 variants, which explained the phenotype of the segregant pool.

**Figure 3.**
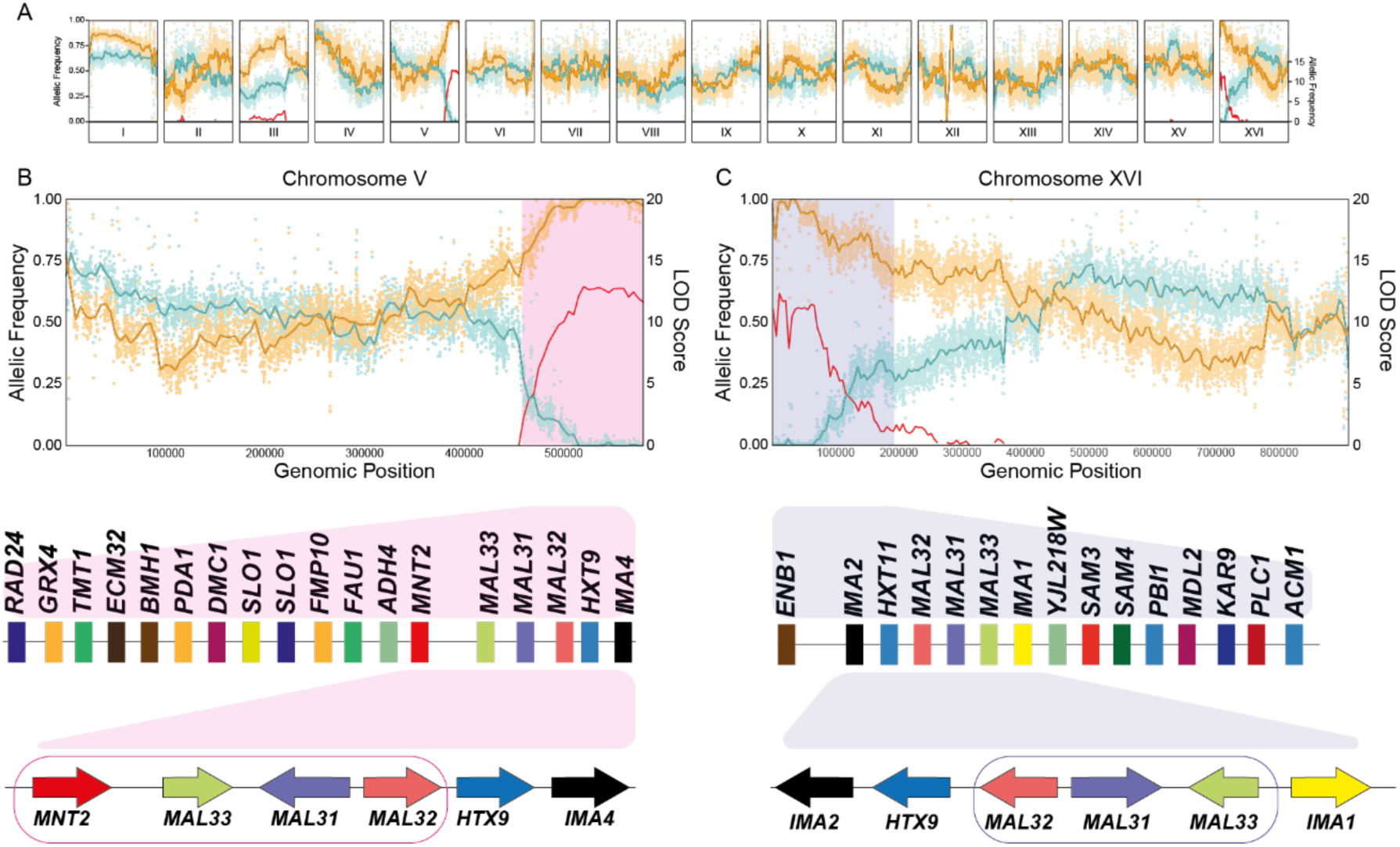
QTL Mapping results in a QC18 x CL467.1 recombinant population. **A)** Allele frequency changes across the genome for the Inferior Pool and Superior Pool (Turquoise and Orange, respectively). LOD Score were plotted against the genome position (Red). Allele frequency of mapped pools against (**B**) Chromosome V-R and (**C**) XVI-L for QTL_1_ and QTL_2_, are represented with pink and blue rectangles, respectively. Candidates’ genes chosen for validation within each QTL are located within the blue and red rectangles, respectively.

Within both QTLs we identified copies of the *IMA* genes (*IMA1*, *IMA2* and *IMA4*), which both encode for isomaltases (**Figure 3B and C**, **Table S7**). In addition, QTL_ChrV_ and QTL_ChrXVI_ contained complete copies of the *MAL* locus, which includes *MAL32* (maltase), *MAL31* (the maltose transporter) and *MAL33* (the transcriptional activator) genes (**Figure 3B and C, Table S8**). Based on this, we selected as candidate QTLs the regions containing the genes of the *MAL* locus for further analysis (**Figure 3B and C**)

### Allelic variation in Chromosome V underlies maltose consumption variation between strains

To determine the phenotypic effect of each QTL in maltose utilization, we performed a reciprocal hemizygosity analysis between QC18 and CL467.1 strains. First, we generated strains lacking either *MAL* locus within each QTL region using a CRISPR-Cas9 mediated strategy (**Figure 4A**) and challenged these mutants in sudden diauxic shift in 20% maltose (**Figure 4B**). Deletion of the *MAL* locus on chromosome V completely abolished growth in 20% maltose in both strains despite the presence of an additional *MAL* copy on chromosome XVI (**Figure 4B**). The inability of one or more genes in chromosome XVI to confer maltose utilization is further supported by mutant analyses: strains lacking the *MAL_ChrXVI_* locus evidenced growth in diauxic shift experiments at 20% maltose. Deletion of this region in CL467.1 resulted in a 0.9-fold reduction in growth compared to the wild type, whereas in QC18, growth increased by 1.2-fold (**Figure 4B**, p-value < 0.05; **Table S9**).

**Figure 4.**
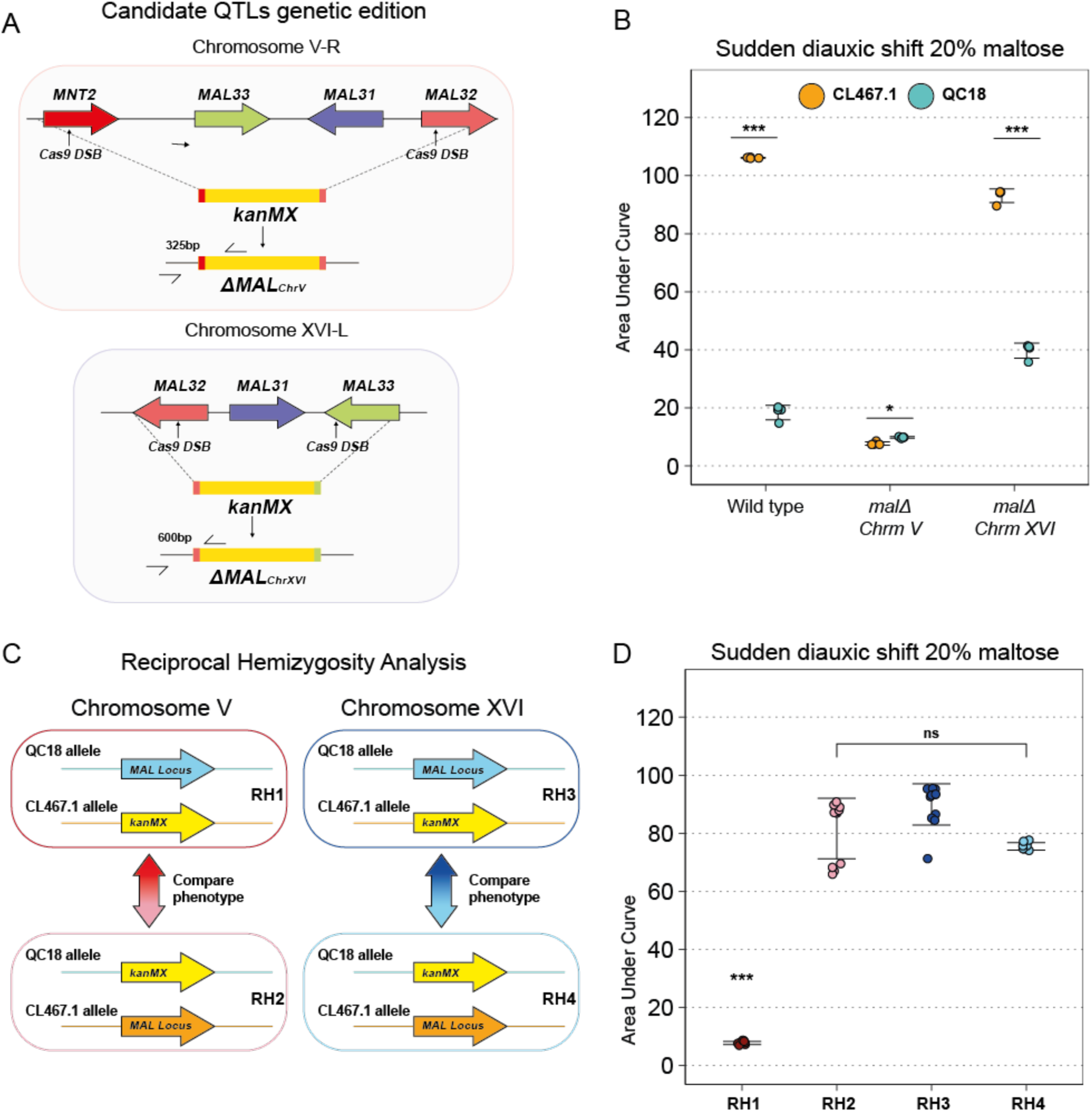
*MAL* locus validation using reciprocal hemizygosity analysis (RHA). **A**) CRISPR-Cas9 editing strategy for candidate QTL knockout. specific guide RNAs targeting each QTL. *MNT2* and *MAL32* targeted for QTL_1_ (Chromosome V-R) and *MAL32* and *MAL33* targeted for QTL_2_ (Chromosome XVI-L). **B**) AUC for QC18 and CL467.1 wild type and mutant strains (*malΔ* for chromosomes V and XVI, respectively), under 20% maltose sudden diauxic shift. **C**) Schematic RHA strain construction. Wild type or mutants QC18 and CL467.1 strains were crossed to generate hemizygous progeny*. E*ach cross represents an isogenic individual that maintains a single functional *MAL_ChrV_* or *MAL_ChrXVI_* allele. **D**) Total growth of reciprocal hemizygote under sudden diauxic shift conditions using 20% maltose as a secondary carbon source. Hemizygotes lacking *MAL_ChrV_* allele are represented as red and pink dots; Hemizygotes lacking *MAL_ChrXVI_* allele are represented as blue and light blue dots. Plotted values correspond to the mean value of eight independent replicates for each strain. The (*) represents different levels of significance between QC18 and CL467.1 strains in one condition (* p < 0.05; ANOVA).

Once the *MAL* locus mutants were constructed in QC18 and CL467.1, crosses were performed between each mutant strain and its corresponding wild-type counterpart (**Figure 4C**). The RHs were analyzed upon sudden shifts in 20% maltose (**Figure 4D**), evaluating their total growth through AUC and compared to both parents and the F_1_ hybrid (**Figure 4D**). RHs carrying the CL467.1 *MAL_ChrV_* locus exhibited an 11.6x greater AUC growth than the opposite RH carrying the QC18 allele (**Figure 4D, Table S10**), demonstrating that allelic differences within the *MAL_ChrV_* locus underlie growth differences under maltose conditions. We observed a similar trend when the same approach was used for the *MAL* locus on chromosome XVI (**Figure 4D**). However, we observed a 1.25-fold growth difference between RHs, with the QC18 *MAL_ChrXVI_* allele exhibiting a lower AUC (**Figure 4D**). These results indicate that allelic variants within *MAL_ChrV_* might exert a more substantial function than *MAL_ChrXVI_*, supporting diauxic shifts under high maltose concentrations.

To identify candidate genes within the two QTL regions, we analyzed the nucleotide sequences of the *MAL* genes from each locus using SNPeff for polymorphism prediction and sequence alignments (**Table S8**). In general, *MAL32* and *MAL31* exhibited a high level of identity across different chromosomes in both strains (> 98%). However, when comparing the *MAL33* loci on different chromosomes, sequence identity dropped below 80%, demonstrating greater sequence divergence than the other two *MAL* genes (**Table S11).** The allelic comparisons between QC18 and CL467.1 revealed high-impact polymorphisms affecting the protein products of *MAL* genes on both chromosomes. However, QC18’s ability to grow on low maltose concentrations suggests that the enzyme encoded by *MAL32* retains functional activity. Interestingly, in *MAL33*, two and one high-impact nucleotide mutations were identified in QC18, leading to a premature stop codon (**Figure 4C**). In contrast, no high-impact mutations were detected in *MAL31*. The lower sequence identity between chromosome copies and premature stop codons in *MAL33* suggests that differences in transcriptional regulation may affect maltose utilization between strains.

### Complementation of MAL33 transcription factor restores maltose utilization in QC18

To validate the contribution of allelic variation in *MAL33*, we conducted a heterologous complementation assay using a multicopy episomal vector (pRS426) to express the different alleles of the *MAL33* locus (Promoter-ORF-terminator). Total growth was assessed upon sudden diauxic shift to 20% maltose. In strain CL467.1, none of the variants ectopically expressing *MAL33* outperformed the wild-type strain (**Figure 5A; Table S12**). However, in QC18, the heterologous expression of *MAL33_ChrV_* alleles from CL467.1 and QC18 resulted in significant growth increases of 2.7-fold and 2.2-fold, respectively, rescuing the maltose consumption phenotype in this strain (*p* < 0.05, ANOVA). Interestingly, expression of *MAL33_ChrXVI_* partly restored the maltose adaptation phenotype only when expressing the QC18 allele, which exhibited a 23% growth increase (**Figure 5B**, *p* < 0.05, ANOVA). However, these growth values remained significantly lower than those obtained with *MAL_ChrV_* variants. These findings suggest that, despite nucleotide-level predictions of impaired functionality, the *MAL33_ChrV_* allele from QC18 is capable of activating downstream maltose-related genes. Thus, the slow growth phenotype may result from regulatory mechanisms or gene dosage effects.

**Figure 5.**
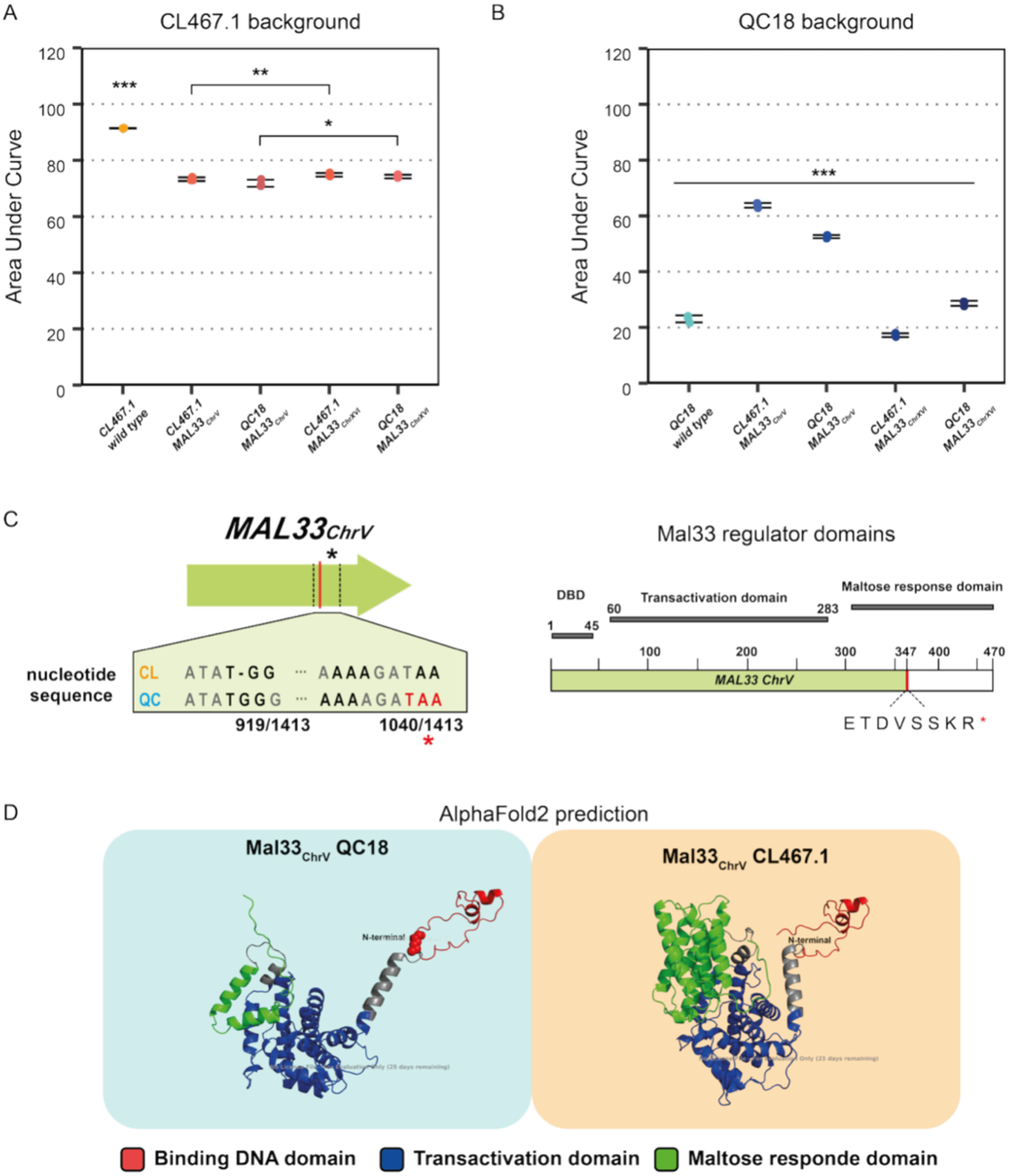
Heterologous complementation assay of the *MAL33* gene. Total growth (AUC) of (**A**) CL467.1 and (**B**) QC18 strains overexpressing *MAL33_ChrV_ or MAL33_ChrXVI_* gene variants. Plotted values correspond to the mean value of three independent replicates for each strain. The (*) represents different levels of significance between QC18 and CL467.1 strains in one condition (* p < 0.05; ANOVA). (**C**) Genetic and functional organization of the *Mal33_ChrV_* activator, with a premature stop codon in the QC18 background, represented as a red asterisk. The domain architecture of the *Mal33_ChrV_* protein is illustrated, highlighting the DNA-binding domain (DBD), transactivation domain, and maltose response domain. (**D**) Predicted 3D structure of the *Mal33_ChrV_* activator protein from QC18 and CL467.1 backgrounds, different domains as the DNA-binding domain (DBD), transactivation domain, and maltose are represented by red, blue, and green.

Based on these results, we analyzed the protein and domain structures encoded by the different *MAL33*_ChrV_ variants. This analysis revealed that QC18 has a truncated protein of 347 amino acids (out of a total of 471), comprising only two out of three major functional domains (**Figure 5C**). The first domain, encompassing a DNA-binding domain containing zinc fingers, is located near the N-terminus at residue 45. The second is a transactivation domain that spans residues 60 to 283. The QC18 Mal33p lacks the third domain, the maltose-responsive activation domain at the protein’s C-terminus. AlphaFold modeling confirmed the truncation in the maltose-responsive domain (**Figure 5C**). Despite this truncation, the transactivation domain retains structural similarity to Mal33p from CL467.1, suggesting that Mal33p from QC18 remains partially functional (**Figure 5D**). However, we found a distinct folding pattern in the DNA-binding domain between alleles. The absence of the maltose-responsive domain and structural differences in the DNA-binding domain likely contribute to the phenotypic differences observed between strains under maltose growth conditions. Overall, these results suggest that allelic variation in *MAL33*_ChrV_, particularly in the maltose-responsive activation domain, underlies the phenotypic differences observed between CL467.1 and QC18 strains.

## DISCUSSION

In this work, we employed a quantitative genetics approach to identify the genetic basis of natural variation in maltose consumption between two Patagonian strains of *S. eubayanus*: QC18 and CL467.1 strains. The QTL analysis identified two loci associated with maltose metabolism, located in the subtelomeric regions of chromosomes V-R and XVI-L. These loci contain complete copies of the *MAL* genes, including *MAL32* (encoding maltase), *MAL31* (maltose transporter), and *MAL33* (transcriptional activator) in both strains (Brouwers et al., 2019). Subtelomeric regions are known to represent hotspots for metabolic gene evolution in yeast and environmental adaptation, often exhibiting high variability and gene content variation that contributes to niche adaptation (Barton et al., 2008; Brown et al., 2010; Cohn et al., 2006; Keeney, 2008; Lambie & Roeder, 1986; Linardopoulou et al., 2005; Rudd et al., 2007; Snoek et al., 2014; Teste et al., 2010; Walther et al., 2014; Yue et al., 2017). In this sense, natural variation in maltose metabolism among natural strains may not be mainly driven by genes previously linked to respiration and glucose repression, such as *HAP5*, *HAP4*, *CIN5*, *PUT3*, *MIG1*, and *SNF1*, since these were not found in our analysis (Molinet et al., 2022; Perez-Samper et al., 2018).

The transition between carbon sources slows down cellular growth for the duration required to activate the expression of genes involved in utilizing alternative sugars (New et al., 2014; Pontes et al., 2024). The diauxic growth of QC18 and CL467.1 was influenced by the type and concentration of the alternative sugar, as well as the diauxic shift approach (sudden or gradual). We observed significant differences in growth kinetics parameters under high maltose concentrations in sudden glucose-maltose conditions, with QC18 exhibiting a substantially reduced ability to grow on 20% maltose compared to CL467.1. In the sudden scenario, the observed lag phase is longer and more heterogeneous compared to the gradual scenario, representing the time required to activate the *MAL* genes (New et al., 2014). Our observations are consistent with previous studies, as the gradual diauxic shift approach from glucose to maltose results in a shorter lag phase under all conditions. In this scenario, the cells are able to detect maltose while glucose is still available, allowing for early adaptation to the diauxic shift, which could be the result of leaky expression of the *MAL* genes (Cerulus et al., 2018; New et al., 2014; Vermeersch et al., 2019). In this sense, the differences between QC18 and CL467.1 may be due to a variable degree of glucose repression. However, this effect was not observed in sucrose or galactose, suggesting a specific regulatory difference in maltose metabolism rather than glucose repression. Sucrose and maltose can be transported by maltose transporters like *MAL31* (Botman et al., 2024). Differences in maltose transport should be reflected in differences in sucrose transport and consumption, which was not the case, suggesting that differences in *MAL32* or *MAL33* could underlie the differences in maltose metabolism.

Polymorphism analysis of *MAL* genes between QC18 and CL467.1 revealed several high-impact mutations in *MAL32* and *MAL33* alleles in QC18, including frameshift mutations leading to premature stop codons. These mutations likely result in truncated maltase and transcriptional activator proteins, consistent with previous studies showing that loss-of-function mutations in *MAL* genes from Holarctic *S. eubayanus* strains severely impair maltose metabolism (Bergin et al., 2022; Brouwers et al., 2019; Peris et al., 2016). Fermentative environments have exerted selective pressure on the *MAL* genes of yeasts such as *S. cerevisiae* and *S. pastorianus*, driving changes in their ability to sense, transport, and metabolize maltose (Gonçalves et al., 2016; Hernández-Vásquez et al., 2024; Peris et al., 2023). This supports that the increase in the copy number of these subtelomeric genes is a feature of domestication and evolution (Brown et al., 2010; Gallone et al., 2016; Ohdate et al., 2018). However, our study provides evidence of how this trait varies across natural isolates unrelated to anthropogenic environments. In this sense, in *Saccharomyces cerevisiae*, each *MAL* locus, despite having the same structure, is genetically unlinked (Needleman & Michels, 1983; Ohdate et al., 2018) and has a different impact on maltose consumption (Day et al., 2002; Gonçalves et al., 2016; Meurer et al., 2017; Nijkamp et al., 2012; Weller et al., 2023). Similarly, we found in *S. eubayanus* that the *MAL* locus, located on chromosome V, makes a greater contribution to the maltose adaptation phenotype than the copy located on chromosome XVI. Our findings suggest that the *MAL* locus located on Chromosome XVI is not sufficient to confer maltose consumption and adaptation ability.

Differences in regulation are a known source of phenotypic variation among strains of the same species (Hernández-Vásquez et al., 2024). An example of this is observed in the laboratory strains W303 and S288C of *S. cerevisiae*, as well as in sake strains, which are unable to consume maltose due to defects in the Mal33 regulatory protein. (Nijkamp et al., 2012; Ohdate et al., 2018). This phenotype is restored through overexpression of the *MAL33* gene from maltose-consuming strains (Brown et al., 2010; Weller et al., 2023). Heterologous expression of the *MAL33_ChrV_* gene from CL467.1 in the QC18 strain restores the maltose adaptation phenotype, confirming that the *MAL31* and *MAL32* genes in QC18, along with their cis-regulatory sequences, are transcriptionally functional. Interestingly, QC18 also restores maltose growth when ectopically expressing its allele of *MAL33_ChrV_*, indicating that the slow maltose consumption would be due to an *MAL33_ChrV_* reduced capacity to activate the *MAL* locus. Protein structure prediction supports an impairment in the maltose-responsive activation domain at the protein’s C-terminal in Mal33p in QC18 (Hu et al., 1999; Ohdate et al., 2018; J. Wang & Needleman, 1996). Consequently, the QC18 *MAL33_ChrV_* gene would be unable to activate the maltose utilization pathway due to a dosage-dependent effect, which is alleviated only by the ectopic expression of these genes. Together with the sequence analysis of the *MAL33_ChrV_* gene, It is possible to infer that this activator has partially lost its functionality and its capacity to respond to intracellular maltose (Change et al., 1988a, 1988b; Gibson et al., 1997), as the gene product still contains the DNA-binding domain and the transactivation domain (Hu et al., 1999; J. Wang & Needleman, 1996). Considering that the maltose sensor has not yet been identified, the maltose-responsive domain of the Mal33p protein represents a candidate responsible for modulating and explaining the differences observed in maltose consumption and utilization. (Horák, 2013; X. Wang et al., 2002).

Mutations in *MAL33_ChrV_* may have arisen recently, as *Quercus robur* is an introduced species in the Southern Hemisphere within the past 500 years and represents an exotic tree within the Patagonian forest (Haneca et al., 2009). Notably, all the known Holarctic *S. eubayanus* strains isolated in the Northern Hemisphere from *Quercus robur* trees have lost their ability to grow and consume maltose (Bergin et al., 2022; Brouwers et al., 2019; Crandall et al., 2023; Peris et al., 2016), suggesting a convergent phenotypic adaptation associated with oak trees. The low ability of QC18 to metabolize maltose might be influenced by the sugar composition of *Q. robur* bark, which primarily consists of glucose and xylose (Miranda et al., 2017). This contrasts with the higher concentrations of complex sugars, such as starch, found in the bark of native tree species like *Nothofagus pumilio* (Molinet et al., 2022). Thus, we hypothesized that *MAL* loci may be essential for survival under *Nothofagus* habitats, unlike the *Quercus* environment, where they may be under purifying selection. In this way, subtelomeric *MAL* gene diversity would represent a reservoir of evolutionary plasticity and a key evolutionary mechanism influencing maltose adaptation among natural *Saccharomyces* isolates. Future research would help characterize the metabolite composition of *Quercus* and *Nothofagus* trees, enabling a deeper understanding of how the host and environment modulate the genomic composition of natural *Saccharomyces* isolates across hemispheres.

## ACKNOWLEDGMENTS

We thank Valentina Abarca for the technical support, PQ acknowledges Hittinger Lab and Biology of Genomes Laboratory for providing infrastructure, laboratory space, and experiment equipment. This research was funded by Agencia Nacional de Investigación y Desarrollo (ANID) FONDECYT program and ANID-Programa Iniciativa Científica Milenio – ICN17_022 and NCN2024_040. FC is supported by FONDECYT grant N° 1220026. PQ is supported by ANID BECAS DOCTORADO NACIONAL Folio 21201057 and Facultad de Química y Biología, Universidad de Santiago de Chile. FMG is supported by FONDECYT POSTDOCTORADO grant N° N°3220597 and PV by FONDECYT INICIACIÓN grant N°11240649. FC, PV and GF are funded by PROGRAMA DE COOPERACIÓN CIENTÍFICA ECOS-ANID ECOS180003 and C23B05. Research in the Hittinger Lab is funded by the National Science Foundation (DEB-2110403), USDA National Institute of Food and Agriculture (Hatch Project 7005101), and in part by the DOE Great Lakes Bioenergy Research Center (DOE BER Office of Science DE–SC0018409) The funders had no role in study design, data collection and interpretation, or the decision to submit the work for publication.

## AUTHOR CONTRIBUTIONS

Conceptualization, P.Q, F.M.G and F.A.C; Methodology, P.Q, F.M.G, N.G, J.G.C and C.M.T; Software P.Q, T.A.P, P.V; Validation, P.Q, F.M.G; Formal analysis, P.Q; F.M.G, G.F and F.A.C; Investigation, P.Q, F.M.G, N.G and C.M.T; Resources, F.A.C, L.F.L, C.T.H and G.F; Data curation, P.Q, F.M.G and F.A.C; Visualization, P.Q and F.M.G; Supervision, F.A.C, L.F.L, C.T.H and G.F; Writing – Original Draft, P.Q, F.M.G, and F.A.C; Writing – review & editing, P.Q, F.M.G, L.F.L, C.T.H, G.F and F.A.C; Funding acquisition, F.A.C and L.F.L Project administration, F.A.C. All authors have read and agreed to the published version of the manuscript.

## SUPLEMENTAL MATERIAL

### FIGURES

**Figure Supplementary 1. Kinetics parameters of *S. eubayanus* strains in sudden diauxic shift experiment using YP supplemented with increasing concentrations of sucrose or galactose (5%, 10%, 20% and 30%). A-B)** Growth kinetics of QC18 (Turquoise) and CL467.1 (Orange) on sucrose. Lag time and Maximum OD, respectively. **C-D)** Growth kinetics of QC18 (Turquoise) and CL467.1 (Orange) on galactose. Lag time and Maximum OD, respectively. Plotted values correspond to the mean value of three independent replicates for each strain. The (*) represents different levels of significance between QC18 and CL467.1 strains in one condition (* p < 0.05; ANOVA).

**Figure Supplementary 2. Kinetics parameters of *S. eubayanus* strains in gradual diauxic shift experiment using YP supplemented with 1% glucose and increasing concentrations of maltose, sucrose and galactose (5%, 10%, 20% and 30%). A)** Growth kinetics of QC18 (Turquoise) and CL467.1 (Orange) on maltose. Total Growth (AUC), Maximum OD and Lag time, respectively. **B-C)** Total growth (AUC) of QC18 (Turquoise) and CL467.1 (Orange) on sucrose and galactose. Plotted values correspond to the mean value of three independent replicates for each strain. The (*) represents different levels of significance between QC18 and CL467.1 strains in one condition (* p < 0.05; ANOVA).

**Figure S3. Bulk Segregant Analysis to QC18xCL467.1 F2 offspring in gradual diauxic shift.** Growth kinetics of QC18, CL467.1, F1 hybrid and F2 segregants in (**A**) glucose 1% and (**B**) gradual diauxic shift experiments, using 20% galactose as secondary carbon source. Parentals strains QC18 and CL467.1 are represented by turquoise and orange lines. Top and bottom 30 F2 segregants were selected from each extreme pool, F2 Inferior and superior segregants are represented by blue and dark orange lines, respectively. Lines show the median OD620nm of three biological replicates

### Tables

**Table S1.** Strains used in this study.

**Table S2.** Growth conditions.

**Table S3.** Primers used in this study.

**Table S4A.** Kinetic parameters in sudden and gradual diauxic shift experiments

**Table S4B. Statistical analysis for AUC, Lag Time and Odmax**

**Table S5.** Area Under Curve for the Bulk Segregant Analysis.

**Table S6.** Genomewide allele frequency in superior and inferior pools.

**Table S7.** Candidate genes in the QTL mapping strategy

**Table S8.** Snpeffect prediction for polymorphisms in QTL candidate regions

**Table S9A.** Candidate QTL knockout strains

**Table S9B. Statistical analysis for Area Under Curve for QTL mutants strains**

**Table S10A.** Reciprocal Hemizygosity Analysis

**Table S10B Statistical analysis for Area Under Curve for RH strains**

**Table S11.** Percentage of identity calculated for the MAL genes

**Table S12A.** Heterologous complementation assay of *MAL33* gene

**Table S12B. Statistical analysis for Area Under Curve in QC18 background**

**Table S12C. Statistical analysis for Area Under Curve in CL467.1 background**

